# The Protein Mat(ters) - Revealing Biologically Relevant Mechanical Contribution of Collagen and Fibronectin Coated Micropatterns

**DOI:** 10.1101/668905

**Authors:** Aron N. Horvath, Claude N. Holenstein, Unai Silvan, Jess G. Snedeker

## Abstract

Understanding cell-material interactions requires accurate characterization of substrate mechanics, which are generally measured by indentation-type atomic force microscopy. Although model extracellular matrix coatings are used to facilitate cell-substrate adhesion, their tensile mechanical properties are generally unknown. In this study a novel tensile mechanical characterization of collagen and fibronectin micropatterned polyacrylamide hydrogels is performed. Our findings reveal that the protein coating itself has measurable and biologically relevant consequences, with ligand-specific tensile resistance of the patterned regions relative to the non-patterned surfaces. To our knowledge our study is the first to uncover a direction-dependent mechanical behavior of the protein coatings and to demonstrate that it affects cellular response relative to substrate mechanics.

## 1. Introduction

Mechanobiology aims to understand how environmental mechanical cues impact cellular activity. Besides the numerous forces to which cells are exposed in their natural microenvironment,^[1]^ one key mechanical signal to which cells respond is the rigidity of the surface to which they adhere.^[2–7]^ This stimulus extensively determines the cellular fate including cell differentiation,^[3,8–10]^ proliferation,^[11,12]^ migration and apoptosis.^[13–17]^ To investigate this environmental rigidity-dependent cell behavior and adaptation, tunable stiffness substrates coated with different extracellular-mimicking ligands have been widely employed.^[18–23]^ Many studies have exploited these platforms to parametrically analyze the phenotypic cellular changes by systematically altering the underlying substrate stiffness and applying different cell-adhesion ligands.^[24–28]^ However, for the interpretation and understanding of the cellular responses, the characterization of the substrate characteristic including its extracellular matrix (ECM) mimicking surface is paramount.

The characterization of such substrates is commonly performed by indentation-type atomic force microscopy (IT-AFM).^[29–31]^ Generally, this is done by indentation with a calibrated cantilever tip, with the measured force-displacement curves fitted to a corresponding indentation model to obtain the elastic modulus of the probed material.^[32–36]^ However, this type of mechano-profiling uses contact models assuming isotropic, homogeneous elasticity which is violated by presence of the physical cues from the protein-layer. Additionally, the estimated stiffness value strongly depends on the indentation depth and the thickness of the protein-layer.^[37]^ Lastly, because of the perpendicular indentation applied this method measures the compressive mechanical properties of the material.

It is known that actomyosin cytoskeleton generated force fluctuations within the focal adhesions mediate ECM-rigidity sensation.^[38]^ In 2D culture conditions cells exert mainly forces parallel to the surface, with out-of-plane forces generally being neglected in traction quantification.^[39]^ With this in mind, we hypothesized that the mechanical properties that cells sense when adhering to soft 2D surfaces could be dramatically affected by the in-plane properties of the deposited ECM coating. To test this, we mechanically characterized fibronectin- and collagen-patterned soft polyacrylamide hydrogels using two different approaches. On one hand we performed standard compression-based AFM micro-indentation using a colloidal particle probe cantilever as indenter and found that differences in stiffness values were minimal between patterned and non-patterned surface regions. In turn, a tensile characterization of identically prepared substrates was done using a biaxial stretcher,^[40]^ revealing significantly higher resistance values of the patterns compared to the surrounding non-patterned surface. Further, we showed that the magnitude of the resistance is protein-specific, with linear elastic behavior of the protein-layer within the tested strain ranges. Finally, we demonstrated that these effects lead to non-negligible artefacts in characterization of tensile cell properties. Specifically, we demonstrate these artefacts in dynamic cell stretching experiments, showing that measures of cellular resistance to an applied tension is biased by the inherent tensile mechanical properties of the protein ligand.

## 2. Results

In the present work, we analyzed the contribution of protein surface-coatings to the bulk mechanical properties of soft hydrogels. For this we developed a new stretchable substrate-preparation method for micropatterned hydrogels to allow the estimation of mechanical properties of patterned and non-patterned regions using compressive (perpendicular to the surface) as well as tensional (parallel to surface) forces (Figure 1). Briefly, an acrylamide-bisacrylamide solution containing fluorescent microspheres was polymerized against protein-micropatterned glass while being centrifuged (Figure 1A). This approach results in a micropatterned polyacrylamide (PAA) hydrogels with encapsulated high density of fluorescent markers beneath their surface (Figure 1B). Although these markers are only needed for the estimation of in-plane deformations, substrates subject to compressive analysis were identically prepared.

**Figure 1:**
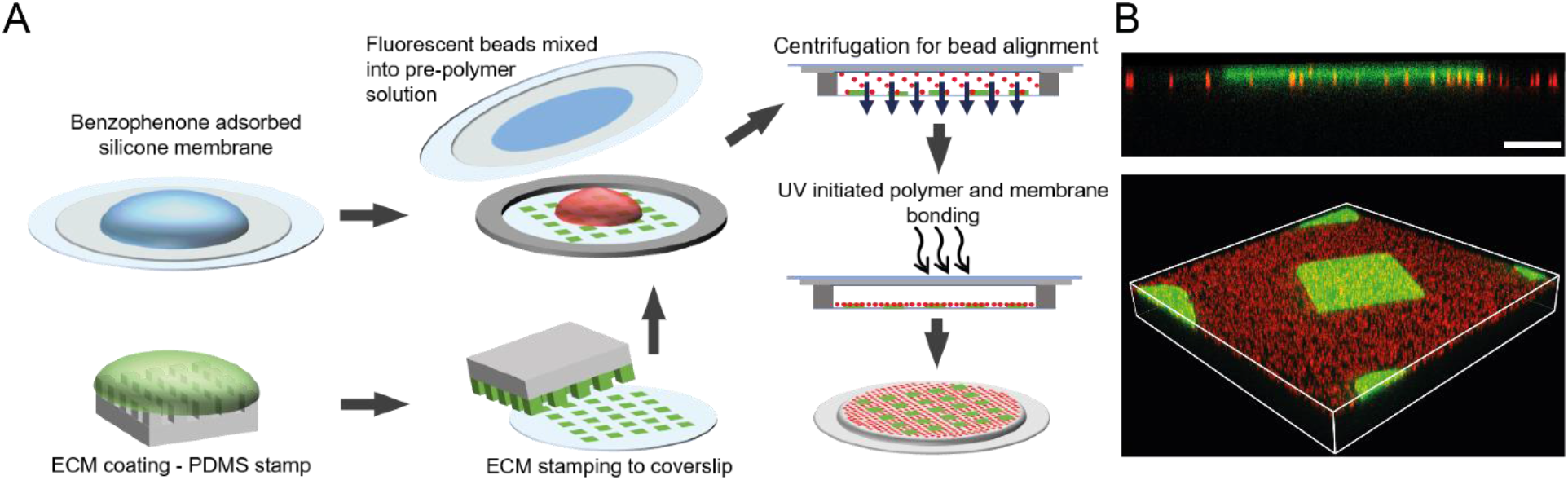
Preparation of stretchable micropatterned substrates with surface aligned beads. (A) Schematic illustration of the preparation method. Benzophenone is adsorbed on silicone membrane, parallel to stamping features to a clean coverslip with a protein pre-coated PDMS stamps. The prepolymer polyacrylamide solution with fluorescent beads is pipetted between the benzophenone-treated silicone membrane and a patterned glass coverslip with a spacer ring. Then this “polymerization sandwich” is centrifuged to align the fluorescent beads to the patterned surface. Next, hydrogel bonding to the membrane is initiated using UV light and after fully polymerization of the gels, the coverslip is carefully removed. (B) Confocal imaging of the micropatterned substrates was used to confirm protein transfer and localization of the fluorescent beads in the upper surface of the substrates (upper panel: lateral view; bottom panel: 3D reconstruction; red – fluorescent markers, green – collagen). Scale bar represents 10 μm.

### 2.1. Compressive stiffness of micropatterned hydrogels

Compressive mechanical analysis of patterned substrates was done using IT-AFM on top of an inverted widefield microscope. As a probe, a spherical indenter of 5 μm diameter was used to obtain force-distance curves, which were interpreted by the Hertz model. The obtained results revealed no statistical differences in the Young’s modulus of fibronectin-patterns and surrounding regions, with recorded values of 7.63 +/− 0.63 kPa and 7.50 +/− 0.48 kPa, respectively (Figure 2). In case of collagen-coated substrates, patterned regions were slightly (~10%) softer than the surrounding substrate, perhaps due to the thickness of the collagen layer. Specifically, the obtained values were 6.31 kPa +/− 0.26 for the collagen-patterns and 7.08 kPa +/− 0.48 in the surrounding regions (p<0.0001) (Figure 2).

**Figure 2:**
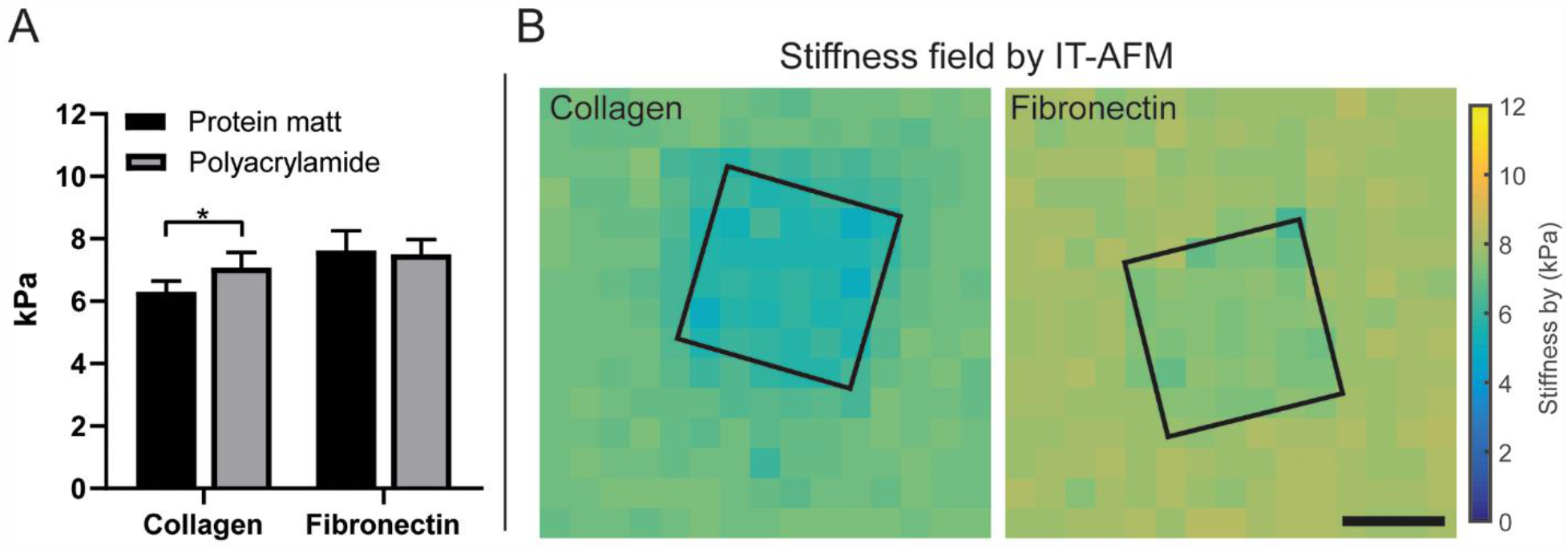
Compressive mechanical characterization of the micropatterned polyacrylamide substrates using IT-AFM. (A) Elastic modulus of the micropatterned polyacrylamide hydrogels, indentations on collagen-patterns (mean= 6.31 +/− 0.34 kPa), fibronectin-patterns (mean= 7.63 +/− 0.63 kPa), or between collagen-patterns (mean= 7.08 +/− 0.48 kPa) and between the fibronectin-patterns (mean= 7.50 +/− 0.48 kPa), n=20, (significance indicated by *, p<0.0001), patterns/group from four different experiment. (B) Example stiffness field of regions with collagen (left panel) and fibronectin (right panel) measured by IT-AFM, square indicates the pattern position; scalebar 20 μm;

### 2.2. Tensile characterization of the micropatterns uncovers significant mechanical differences

For the measurement of the tensile properties of the protein patterns we used a previously developed system based on a commercially available pressure-actuated biaxial stretcher (StageFlexer) built around an upright fluorescent microscope.^[40]^ Substrate drift caused by the biaxial deformation was corrected using fiducial fluorescent markers and a computer-assisted tracking and stage control system. Using this platform, we first calculated the surface strains between minimal actuation (10 kPA applied actuator vacuum) and maximal actuation (60 kPa vacuum) from the displacement of the fluorescent beads compared to the non-stretched, initial state of the membrane (Figure 3B,C). We observed that the average surface strain and the strain below the patterns (Figure 3A,C) increased linearly in the studied range (Figure 4A,B). In addition, independently of the immobilized ECM protein, the strain under the patterns was smaller than the average surface strain. The mean principal strain drop (MPSD) is a metric introduced to quantify the resistance of the patterns to the applied strain and can be defined as

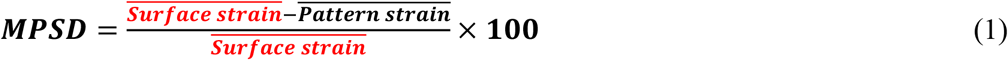

**Figure 3:**
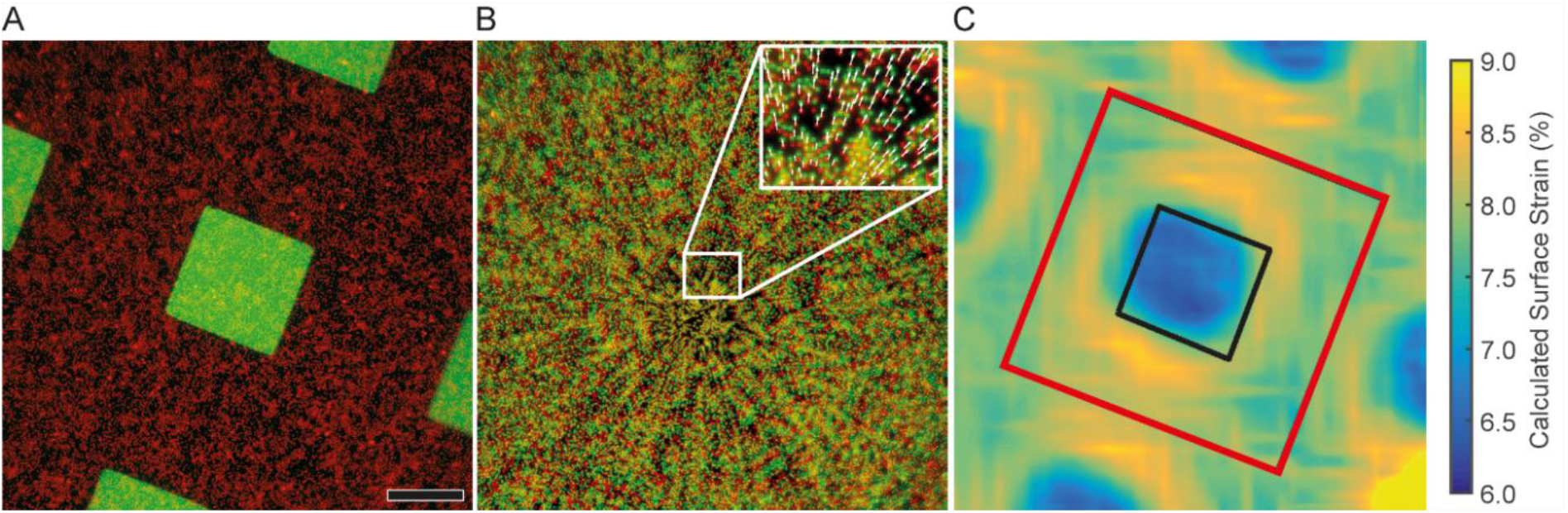
Tensile characterization of the patterned substrates. (A) Pseudo colored example image of a patterned substrate in relaxed state (red – fluorescent labelled microspheres; green – protein mat). (B) Images of pre-(red) and post-stretched (green) bead images were used to determine the displacement field using image processing feature detection. (C) Reconstructed surface strain was calculated from the displacement field. The black and red outline defined areas are used to calculate the average pattern and surface strain, respectively. Scale bar 20 μm.

In case of the collagen-patterns the MPSD slowly decreased linearly from 10.51 +/− 2.71 (10 kPa applied vacuum, approximately 1.67% substrate strain) to 9.75 +/− 2.61 (60 kPa applied vacuum, or approximately 10% substrate strain), being however no statistically significant differences between any of the measured MPSDs. For fibronectin-patterns the calculated MPSD was significantly smaller (1.90% +/− 1.26) and did not show any trend or statistically significant differences in the range of the analyzed strains (Figure 4C).

**Figure 4:**
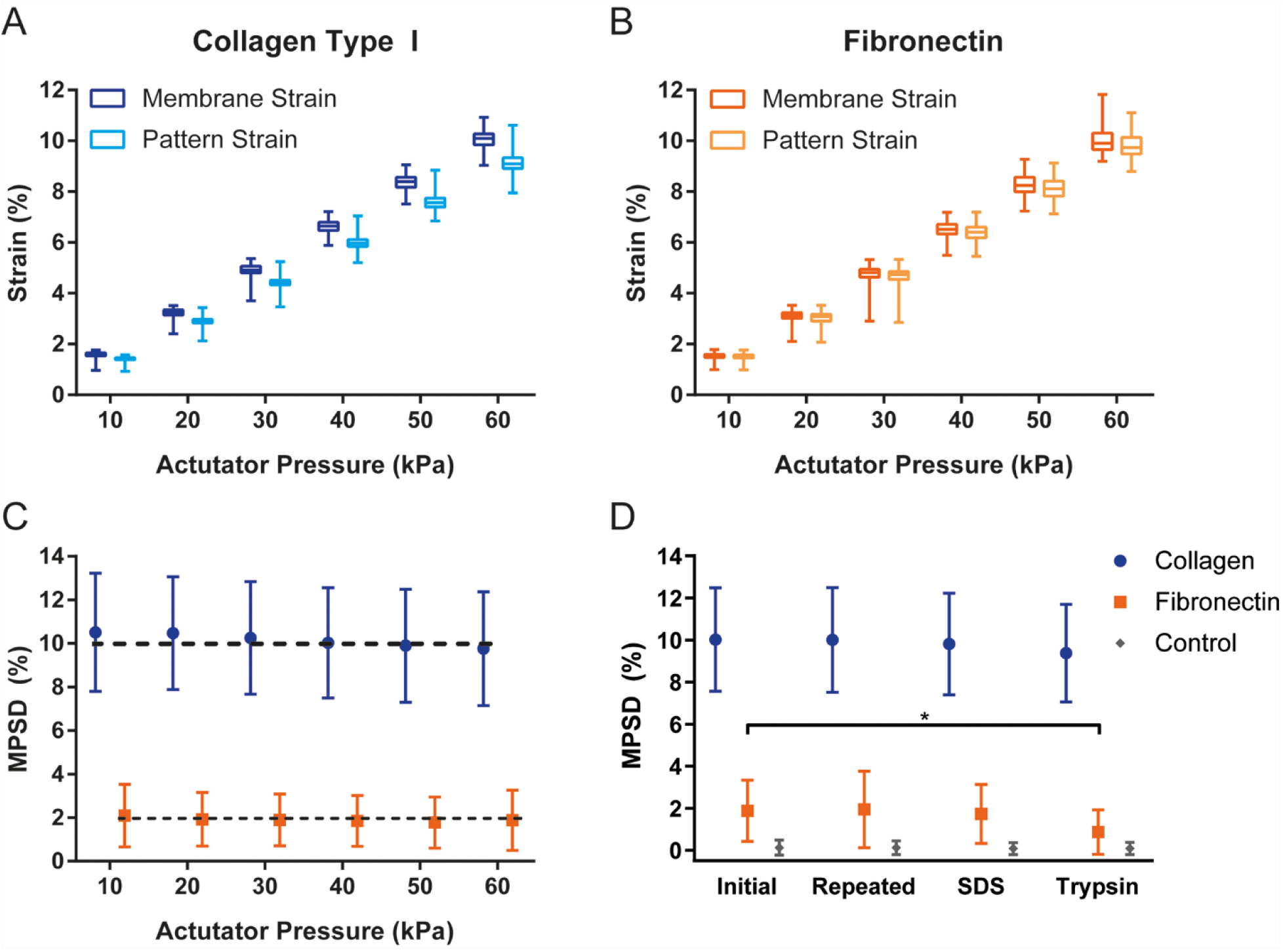
Linearity, durability and bio and chemical treatment of the protein mats. (A,B) The measured change of membrane and pattern strains show linear dependence of the applied pressure range. (C) Calculated MPSD values are constant over the measured actuator pressure range showing the linear elastic behavior of the protein mat. At 8% strain the collagen micropatterns show a significantly higher resistance value to biaxial stretch (MPSD ~10%), than the fibronectin micropatterns (MPSD ~2%). (D) Cyclic loading (up to 50 cycles) does not lead to changes in the mechanical properties of the protein islands. SDS does not change the mechanical behavior of the micropatterns. However, trypsin causes significant drop of the resistance the fibronectin-micropatterns; (significance indicated by *, p<0.001)

To determine the mechanical stability of the protein mat we quantified the MPSD values before and after the application of a cyclic loading protocol consisting of 50 cycles at 0.1 Hz with the pressure-actuator set to 50 kPa (approximately 8.33% biaxial strain). Both, collagen and fibronectin protein mats did not display fatigue and in both cases the MPSD remained stable at 10.01% +/− 2.46 for collagen and 1.91% +/− 1.63 for fibronectin-patterns (Figure 4D). Because TFM is one of the techniques in which the hereby described tensile properties of ECM protein coatings are central, we further analyzed the impact of the cell-dissociation solutions commonly used in such experiments on the mechanical stability of the patterns. In these experiments after the initial MPSD calculation, we incubated the hydrogels in either 0.1% sodium dodecyl sulphate (SDS) or in a commercially available 5% trypsin solution before mechanically testing the membranes again. In both pattern-types a similar behavior was observed with only minor changes in the MPSD before and after the treatment (Figure 4D).

### 2.3. Artefactual impact of the mat on the measured tensile resistance of adherent cells

Finally, we investigated the impact of the tensile properties of the protein mats on cell experiments. For that we allowed 3T3 fibroblasts to adhere to the patterns for 6 hours and quantified the combined MPSD caused by the cells and underlying patterns (MPSD^*cell+pattern*^; Figure 5A, left panels) using the same method as before. Next, we dissociated the cells by incubating the hydrogels in 0.1% SDS and calculated the remaining MPSDs (MPSD^*pattern*^). As it was the case in previous experiments, the MPSD^*pattern*^ of fibronectin was significantly smaller than that of collagen-patterns (1.74% +/− 1.66 vs. 5.03% +/− 1.65 respectively), (middle panels in Figure 5A). By calculating the difference between MPSD^*cell+pattern*^ and MPSD^*pattern*^ we estimated the actual resistance of the cell to the substrate deformation (MPSD^*cell*^), without the mechanical effects of the pattern (Figure 5A, right panels). As example in the typical outcome shown in Figure 5A the calculated MPSD^*pattern+cell*^ was higher for the cell on collagen (collagen 13.33% +/− 4.31 vs. fibronectin 12.56% +/− 6.51), but, due to the lower tensile stiffness of the fibronectin mat, the corrected MPSD^*cell*^ values follow the opposite trend with the cell sitting on fibronectin being stiffer. Specifically, MPSD^*cell*^ for the cell sitting on collagen (example Figure 5B) was 8.30% +/− 4.47 and that on the fibronectin micropattern was 10.82% +/− 5.46 (Figure 5C).

**Figure 5:**
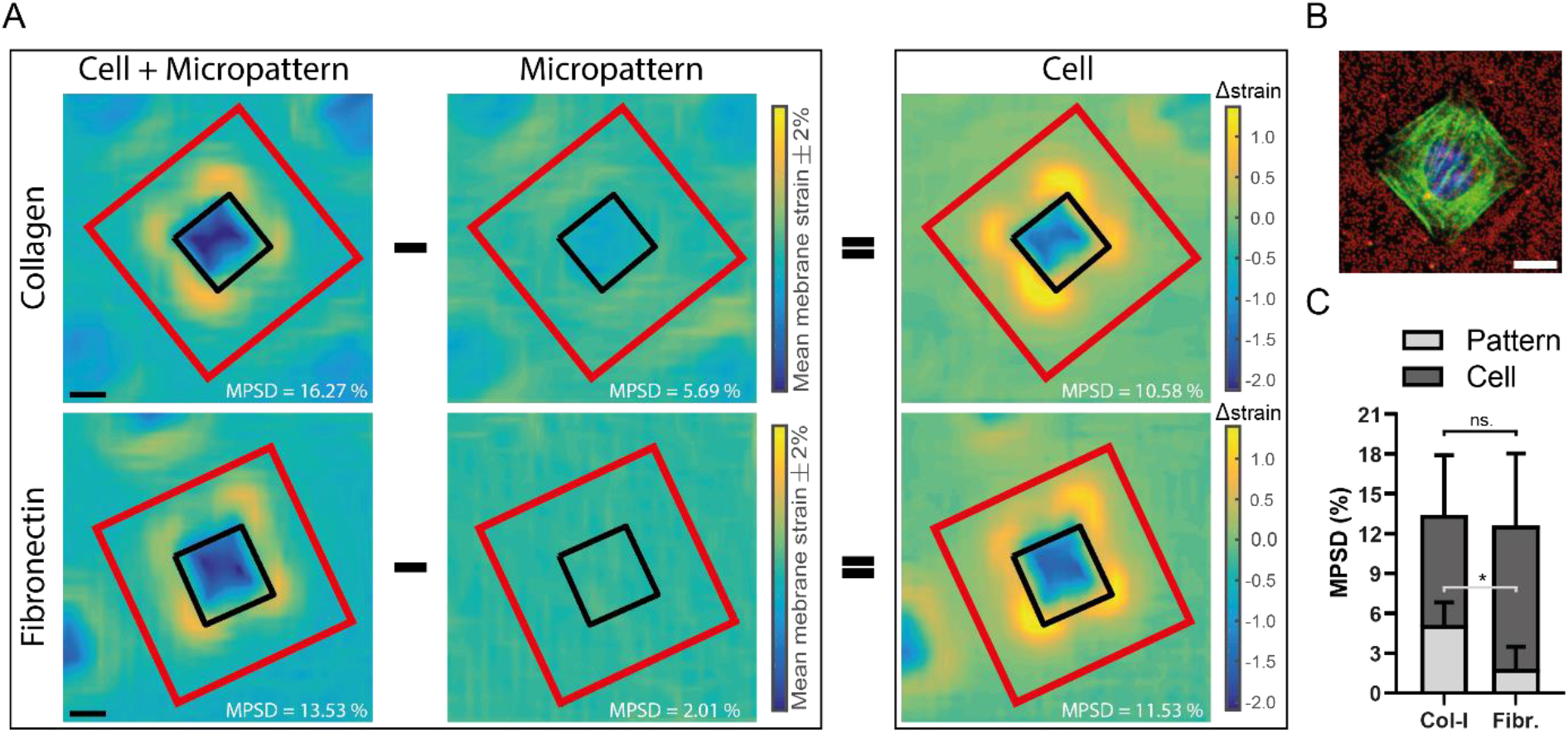
Quantification of the impact of the mechanical properties of the deposited ECM on the estimation of cell tensile stiffness. (A) Strain maps corresponding to collagen (upper row) and fibronectin (bottom row) micropatterns with adhering 3T3 cells (left panels) and after SDS treatment of the cells (middle panels). As the apparent cellular tensile stiffness is biased by the underlying protein mat, to calculate the actual MPSD of the cell the strain drop caused by the pattern is subtracted (right panels). Scale bar 20 μm. (B) Pseudo colored example for a morphology controlled 3T3 fibroblast cells adhered to a 35×35 μm^2^ collagen-coated pattern (red – fluorescent labelled microspheres; green – actin network, blue – nucleus). Scale bar 10 μm. (C) Although cells adhered to collagen micropatterns initially appear stiffer (higher resistance to externally applied strains) than the cells on fibronectin patterns, subtraction of the contribution of the pattern reveals opposite cellular trends, highlighting the artefact imposed by the collagen-mat. Statistics: *, p<0.0001, n.s., not significant

## 3. Discussion

Mechanobiology experiments frequently make use of elastic surfaces onto which extracellular matrix (ECM) proteins, in most cases collagen or fibronectin, are immobilized to promote cell-adhesion.^[30,41–45]^ Because substrate rigidity is a major mechanical regulator an accurate measure of substrate stiffness is often central for the correct interpretation of biological response.^[2,3,7]^ For that, mechanical tests of the substrates are commonly performed, with indentation-type atomic force microscopy (IT-AFM) being the most widely used technique.^[29,30,46]^ Nevertheless, such compressive analysis are insensitive to inherent tensional rigidity of the assemblies formed after protein immobilization.^[47,48]^ With this in mind, we established an approach for the generation of micropatterned polyacrylamide (PAA) hydrogels that enables the comparison of the compressive and tensile properties of ECM-coated and contiguous non-coated regions. Using this platform, we found that although minimal differences were detected in the compressive properties of the non-patterned and patterned regions, substantial differences existed when considering tensile strains in the patterned regions, which were dramatically smaller than in the non-coated regions of the bulk-substrate. In addition, these mechanical properties were found to be protein-specific, with collagen-patterns displaying larger resistance to deformation than fibronectin coated patterns. The ECM islands of both ligands showed high mechanical stability, retaining their mechanical properties after cyclic loading and incubation with cell-dissociation solutions.

Previous studies have extensively focused on revealing changes in cell phenotype as dependent on ECM composition and the mechanical properties of the underlying substrate. On one hand, changes in composition can fundamentally alter cellular response to substrate stiffness through the involvement of differential ECM receptor activity.^[24,26,49]^ On the other hand, stiffening of the mechanical properties or orientation of the ECM can drive cell phenotype without major changes in the cell integrin ligation profile.^[50–52]^ The tensile properties of deposited ECM proteins exposed by our study have potential relevance in experiments focusing on mechanotransduction. For instance, in 2D traction force microscopy (TFM) cell-generated traction forces are inferred from the substrate deformations in combination with the estimated mechanical properties of the used compliant material.^[53,54]^ In such experiments, compressive stiffness is a very widely used to characterize the general mechanical properties of a substrate. However, it is essential to note that any disconnect between the compressive elasticity of a substrate and the tensile elasticity of that substrate is potentially highly problematic. Cells cultured on flat surfaces exert predominantly in-plane tractions with minor out-plane (normal) components with the consequence that tensile properties of substrates are substantially more relevant.^[39]^ In this sense, the present study highlights a potentially major source of experimental artefact that appears to have gone unappreciated until now.

The present work also highlights the potential necessity to account for active cell remodeling of the protein mat on mechano-variant substrates over the course of an experiment. Cells actively remodel their local environment through deposition of newly synthesized proteins, crosslinking, and degradation of the existing ECM,^[55–59]^ with expected changes in mechanical properties over culture time. Because the magnitude and trend of these changes is likely to be affected by the nature of the cells in the various experimental conditions, it is plausible that tensile testing of the substrate mechanics (as opposed to the standard compressive testing that is done) might be essential to a correct interpretation of experimental outcomes. Such a tensile characterization may even represent an improved functional assay for quantifying ECM remodeling processes, such as the ECM stiffening caused by tumor cells.^[57,60]^

Our observations also highlight potentially serious artefactual impact of protein deposition with regard to the widely used method of traction force microscopy. In these experiments the tensile traction estimation relies on the quantification of the cellular resistance to the applied tensile load and therefore the measured tension is highly affected by the underlying protein resulting in deceptive interpretation of the outcome.^[37]^ Use of minimal cell-adhesion peptides that do not self-assemble into larger structures, such as the fibronectin-derived tripeptide Arg-Gly-Asp (RGD)^[61]^ or its collagen equivalent GFOGER triple helical motif,^[46,62]^ could effectively decouple the biophysical and biological aspects of cell-substrate interaction as their mechanical contribution presumably is negligible.

In addition to its potential relevance to static cell culture systems, non-neglectable tensile properties of deposited ECM proteins also have a dramatic impact on dynamic (stretch) mechanical loading of adherent cells.^[40]^ For instance, we demonstrate that use of collagen coated substrates is likely to diminish transfer of applied substrate strains to the cell when compared to fibronectin. In future work, the strain distribution of fibers within the protein mat may itself be of interest interpreting biological outcomes.

Taken together, our results show that due to the different nature and self-assembly properties of ECM proteins their immobilization on soft substrates can cause ligand-specific, direction-dependent mechanical properties, which can then play a determinant role in stiffness-driven cellular responses. Collectively these results emphasize the importance of accurate and direction-specific substrate characterization for reliable control of cellular behavior and biomechanics.

## 4. Conclusion

In the present work we expose a relevant discrepancy between compressive substrate stiffness, generally used as proxy of substrate mechanical properties, and the surface tensile stiffness perceived by cell adhering to them. We introduced a method for quantitative tensile stiffness characterization of the micropatterned protein mat and used it to demonstrate that collagen and fibronectin coatings have inherent tensile properties that affect baseline physical properties of the substrate surface. This inherent tensile property of the protein mat potentially plays a crucial role in stiffness-dependent cell behavior. In addition, we demonstrated that protein-specific tensile properties can translate to systematically biased biological outcomes and misleading conclusions from experimental data. These results indicate that the uncovered direction-specific mechanical property of protein coatings has potentially critical implications for research focused on cell-material interactions.

## 5. Experimental Section

### Design and production of protein-coated PDMS stamps

SolidWorks software was used to design a chromium mask with 1225 μm^2^ square features, which was then fabricated via standard soft lithography technique at the Center of MicroNano Technology (Lausanne, Switzerland) using a Heidelberg VPG200, Photoresist LASER Writer. The positive photoresist (AZ ECI 3000) spin-coated silicon wafers were exposed to UV light and developed with ACS 200 Gen 3. The negative silicon wafers were silanized overnight in a vacuum desiccator before molding 10:1 PDMS (Sylgard^®^ 184, Dow Corning DC 184) on them. After curing PDMS at 70°C overnight stamps were sonicated in ethanol 15 minutes then washed with water and blow dried with nitrogen. The PDMS stamps were inked either with 100 μg/ml 1:1 ratio of rhodamine-conjugated fibronectin (Cytoskeleton FNR01) and non-labelled fibronectin (Sigma-Aldrich F2006), or 100 μg/ml 1:1 ratio of FITC conjugated collagen (Anaspec AS-85111) and non-labelled collagen (Corning 354236) and for 1 hr. The stamps were blow dried with nitrogen and placed in conformal contact with previously ethanol and water cleaned 30 mm glass coverslips for 1 minute.

### Preparation of polyacrylamide hydrogels with defined mechanical properties

Micropatterned polyacrylamide gels on silicone membranes were prepared with an integration of previously described protocols (Figure 1A).^[63,64]^ Briefly, 50 mm (*∅*) round glass coverslips were adhered to the bottom of the 47 mm (*∅*) silicone membranes (0.5 mm thick for stretching and 0.125 mm for IT-AFM, Specialty Manufacturing, Saginaw) to prevent environmental oxygen to diffuse through the membrane and therefore inhibit polyacrylamide (PAA) polymerization. Next, silicone membranes were incubated in 10% (w/v) benzophenone dissolved in water/acetone mixture (35:65 w/w) for 60 seconds, then rinsed with methanol two times before placing them a vacuum desiccator for 30 minutes. The previously microstamped coverslips were placed into a custom fabricated metal holder and fixed them in place using a 180 μm thick spacer ring (14 mm inner *∅*, 34 mm outer *∅*) on top of them. A solution containing 10/0.13 (w/v) acrylamide/bisacrylamide (Sigma-Aldrich A4058/M1533) and 0.02% 0.5 μm fluorescent beads (Invitrogen™, FluoSpheres™, F8812, F8813) in water and degassed for 30 minutes. 0.0005% v/v tetramethylethylenediamine (BioRad) was mixed into the prepolymer solution and before adding 0.005% w/v acrylic acid N-hydroxysuccinimide (previously dissolved in DMSO, Sigma-Aldrich), pH of the solution was neutralized with HCl (0.0054 % 1 M) to avoid the dissociation of the ester group. Immediately after the addition of ammonium persulfate (BioRad, 0.02% w/v) 30 μl prepolymer solution were pipetted onto the micro-stamped coverslips inside of the spacer. Silicone membranes were vented to nitrogen and then placed on top of the prepolymer with the functionalized side facing the solution. In order to align the beads close to the gel surface, this assembled unit was centrifuged in a swinging bucket centrifuge for 15 minutes at 4200 g. Next, the adsorbed photoinitiator was UV activated by immediately spacing the gels into a UVO Cleaner (Jelight Model 42) at an approximate distance of 1 cm from the UV lamp (20 mW). After polymerization, coverslips were separated from the gels under phosphate-buffer saline (PBS) and patterned hydrogels were incubated in BSA (5% (w/v) in PBS) for 1 hour at room temperature. After passivation, the gels were washed three times with PBS and kept under PBS solution at 4°C until use.

### Indentation-type atomic force microscopy (IT-AFM)

IT-AFM was performed using a Flex-ANA (Nanosurf, Basel) build on top of a widefield microscope (iMic, TILL Photonics, Thermo) equipped with a 20X 0.7 N.A. objective and 10 μm diameter borosilicate glass ball (sQUBE, CP-qp-CONT-BSG, 0.1 N/m) modified cantilevers. The spring constant of each cantilever was determined by thermal tuning and the deflection sensitivity was calibrated by indentation against glass surface. 75×75 μm^2^ areas including 35×35 μm^2^ protein patterns were measured with an indentation spacing of 6 μm.

Five randomly selected areas per substrates of four independent sample per protein ligand were characterized. To calculate the Young’s modulus (E), Hertz model with spherical indenter was fitted to the recorded indentation force curves using custom MATLAB scripts up to 500 nm indentation depth. Based on the recorded indentations positions, the indentations were separated pattern and non-pattern regions. From every recorded indentation field, average values for the pattern and the non-pattern regions were calculated and these values were analyzed across the measured different fields and samples.

### Tensile characterization

For tensile testing a custom imaging platform build on a Leica DM5500 upright microscope equipped with a 40X, 0.8 N.A. water immersion objective was used^<bartalena2011>^. The mechanical resistance of the different protein micropatterns were either linearly tested (0-10%) or evaluated in cyclically loaded fashion (8% strain, 0.1 Hz). The stretching protocols were executed via a pressure-actuated biaxial stretcher (StageFlexer) at 37°C. To test the potential effect of SDS or trypsin incubation on the pattern integrity, the patterns were treated with 0.1% sodium dodecyl sulfate (SDS) solution (in PBS) or 5% Trypsin (in PBS). The surface strain (deformation beneath and surrounding the pattern) was determined by tracking fluorescent beads on the surface using a custom developed MATLAB algorithm. The normalized mean principal strain drop (MPSD) was defined as the difference between the average surface strain and mean strain beneath the pattern, divided by the average surface strain (Figure 1D). For the protein mat characterization, 15 different patterns per each at least three independent samples were analyzed in each ligand group. For the cell experiments, 32 cells on collagen-pattern and 11 cells on fibronectin-pattern were analyzed from at least three independent experiments.

### Cell culture & cell experiment

NIH/3T3 cells were cultured at 37°C, 5% CO2, in DMEM F12 (Sigma,7002211), supplemented with 10% FBS and with 1% Penicillin/Streptomycin. For the mechanical testing, 65 cells/μm^2^ were seeded on the hydrogels and allowed to adhere to the patterns for 6 hours. Two hours before the experiment the cell cytoskeleton was stained by the addition of the live-dye SiR-actin (SpiroChrome) at a final concentration of 500 nM.

### Confocal imaging

Confocal stacks of the substrates were performed with a Nikon A1plus confocal microscope equipped with a Plan APO 60X, 1.40 N.A. oil immersion objective. Used excitation/emission wavelengths for the two fluorescent channels: 488/525nm and 561/595nm. Image stacks were acquired at 0.125 μm intervals for a total distance of 15 μm. Z-stacks were analyzed and exported using the NIS-Elements Viewer.

## Acknowledgements

We thank the Swiss Center for Musculoskeletal Biobanking (SCMB) for granting us access to their Nikon Confocal Microscope. We also gratefully acknowledge the support provided by Christian Bippes, NanoSurf. A.N.H. thanks for JSPS fellowship received to conduct research in Japan. This work was supported by the Swiss National Science Foundation (grant numbers 165670, 138221, 118036 to J.G.S.)

## Table of contents

Cell adhesion to synthetic hydrogels is facilitated by coating of the substrates with extracellular proteins. Nevertheless, the mechanical contribution of these proteins is generally neglected and only the bulk properties of the material are considered. Herein, collagen and fibronectin micropatterned polyacrylamide hydrogels are mechanically characterized revealing minimal compressive, but significant tensile properties of the protein-patterns with demonstrated artefactual impact on cell mechanics studies.

## ToC figure

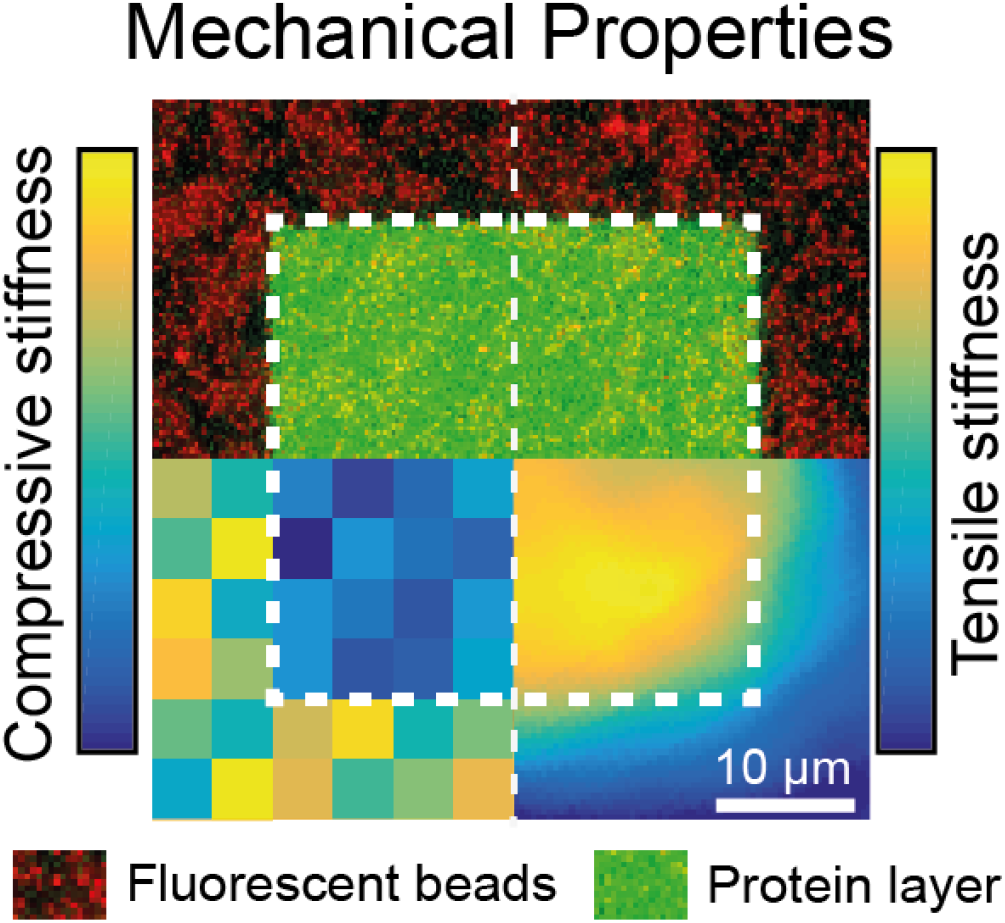

